# Zika virus induces mitotic catastrophe in human neural progenitors by triggering unscheduled mitotic entry in the presence of DNA damage while functionally depleting nuclear PNKP

**DOI:** 10.1101/2021.08.27.458001

**Authors:** Malgorzata Rychlowska, Abigail Agyapong, Michael Weinfeld, Luis M. Schang

## Abstract

Vertical transmission of Zika virus (ZIKV) leads with high frequency to congenital ZIKV syndrome (CZS), whose worst outcome is microcephaly. However, the mechanisms of congenital ZIKV neurodevelopmental pathologies, including direct cytotoxicity to neural progenitor cells (NPC), placental insufficiency, and immune responses, remain incompletely understood. At the cellular level, microcephaly typically results from death or insufficient proliferation of NPC or cortical neurons. NPCs replicate fast, requiring efficient DNA damage responses to ensure genome stability. Like congenital ZIKV infection, mutations in the polynucleotide 5’-kinase 3’-phosphatase (PNKP) gene, which encodes a critical DNA damage repair enzyme, results in recessive syndromes often characterized by congenital microcephaly with seizures (MCSZ). We thus tested whether there were any links between ZIKV and PNKP.

Here we show that two PNKP phosphatase inhibitors or PNKP knockout inhibited ZIKV replication. PNKP relocalized from the nucleus to the cytoplasm in infected cells, co-localizing with the marker of ZIKV replication factories (RF) NS1 and resulting in functional nuclear PNKP depletion. Although infected NPC accumulated DNA damage, they failed to activate the DNA damage checkpoint kinases Chk1 and Chk2. ZIKV also induced activation of cytoplasmic CycA/CDK1 complexes, which trigger unscheduled mitotic entry. Inhibition of CDK1 activity inhibited ZIKV replication and the formation of RF, supporting a role of cytoplasmic CycA/CDK1 in RF morphogenesis. In brief, ZIKV infection induces mitotic catastrophe resulting from unscheduled mitotic entry in the presence of DNA damage. PNKP and CycA/CDK1 are thus host factors participating in ZIKV replication in NPC, and pathogenesis to neural progenitor cells.

**Significance:** The 2015-2017 Zika virus (ZIKV) outbreak in Brazil and subsequent international epidemic revealed the strong association between ZIKV infection and congenital malformations, mostly neurodevelopmental defects up to microcephaly. The scale and global expansion of the epidemic, the new ZIKV outbreaks (Kerala state, India, 2021), and the potential burden of future ones pose a serious ongoing risk. However, the cellular and molecular mechanisms resulting in microcephaly remain incompletely understood. Here we show that ZIKV infection of neuronal progenitor cells results in cytoplasmic sequestration of an essential DNA repair protein itself associated with microcephaly, with the consequent accumulation of DNA damage, together with an unscheduled activation of cytoplasmic CDK1/Cyclin A complexes in the presence of DNA damage. These alterations result in mitotic catastrophe of neuronal progenitors, which would lead to a depletion of cortical neurons during development.

## Introduction

Since its identification in 1947, Zika virus (ZIKV) had been associated with mostly mild and asymptomatic infections until the recent international epidemic with 245,000 confirmed, and 820,000 suspected, cases (1). It took 62 years from the first documented human infection in 1954 (2) to this international epidemic in 2015-2018 to make evident the association of ZIKV infection with the congenital neurodevelopmental pathologies currently referred to as a congenital ZIKV syndrome (CZS) (3). The most significant burden of CZS is congenital microcephaly. Infants born with microcephaly have a range of health problems and suffer lifetime consequences (4, 5). Despite the global health burden, relatively little is yet known about the mechanisms of ZIKV-induced microcephaly.

ZIKV is a unique vector-born flavivirus in that it has two additional routes of transmission, sexual (6–8) and vertical (9–13). The risk of vertical transmission is as high as 46% in the first trimester of pregnancy, and vertically transmitted ZIKV can infect the developing fetal brain producing CZS in about 9% of babies born to mothers infected in the first trimester (14). Neither the mechanism of vertical or sexual transmission, nor the tropism for the developing fetal brain or the microcephaly mechanisms are fully understood. The fetal neurotoxicity of ZIKV is likely the result of a combination of multiple factors, including placental insufficiency and inflammatory responses, activation of innate immunity, and direct ZIKV induced cytotoxicity to neural progenitor cells (recently reviewed in (15, 16)). At the cellular level, congenital microcephaly is caused by the depletion of the neural progenitor cells (NPC), which are responsible for brain development, either by cell death or premature differentiation (17). Proliferating NPC are the primary target for ZIKV, as demonstrated in human autopsies, animal models, and in vitro cell culture systems. ZIKV infection of NPC leads to reduced proliferation and cell death (reviewed in (16, 18)).

Zika virus (ZIKV) is a mosquito-born RNA virus in the *Flavivirus* genus of the *Flaviviridae* family, which comprises several other important arthropod-borne human pathogens like dengue virus (DENV), yellow fever virus (YFV), West Nile virus (WNV), and Japanese encephalitis virus (JEV). Like all viruses in this family, ZIKV has a single stranded, positive sense RNA (ss(+)RNA) genome encoding a single ORF, which is directly translated into a precursor polyprotein (19). The polyprotein is then proteolytically cleaved into the individual viral proteins, including the structural proteins, capsid, E and prM/M, which are incorporated into viral particles, and the non-structural (NS) proteins, NS1, NS2A, NS2B, NS3, NS4A, NS4B and NS5, which are expressed in the infected cell and responsible for viral replication and assembly (20, 21).

Like all ss(+)RNA viruses, ZIKV has a cytoplasmic replication cycle and remodels the endoplasmic reticulum (ER) membranes to assemble the viral replication factories (RF) (22). The characteristic ER rearrangements in flavivirus infections take the shape of dilated ER cisternae containing numerous invaginated vesicles (Ve), vesicular packets (Vp), tubular network of convoluted membranes (CM), and zippered ER (zER) (22–26). These RF enable the concentration of the viral factors required for replication and assembly of progeny virions. Invaginated towards the ER lumen with a narrow opening to the cytoplasm, the Ve contain fully assembled viral replication complexes (RC). Such a semi-open structure is crucial for recruiting the cytoplasmic viral enzymes NS3 (protease and helicase) and NS5 (methyltransferase and RNA-dependent RNA polymerase), while at the same time shielding the viral genomes from recognition and degradation (22, 27). The formation of RF is thus essential for the viral replication cycle.

Several cellular and viral factors play important roles in the biogenesis of RF (recently reviewed in (28)). One of them is the viral non-structural protein 1 (NS1), which localizes to the ER lumen and binds to the inner leaflet of the ER membrane via hydrophobic interactions. Free luminal NS1 also enters the secretory pathway and is secreted (29, 30). Membrane binding and dimerization or multimerization of NS1 induces membrane curvature leading to the formation of invaginations directed towards the ER lumen, thus initiating the shaping of the Ve (25). There is also mounting evidence that flaviviral NS2A, 2B, 4A and 4B, as well as several cellular proteins, play important roles in the biogenesis of RF (28). The formation and maintenance of ZIKV RFs is thus a combined effort of several NS and cellular proteins, not all of which have been identified.

ZIKV is only a minor cause of global microcephaly, with over 3,720 confirmed CZS cases worldwide (1). The vast majority of microcephalies with identified etiology result from genetic disorders, including mutations in one of the essential DNA damage repair proteins, polynucleotide kinase 3’-phosphatase (PNKP). These mutations are associated with a primary form of microcephaly named microcephaly, seizures, and developmental delay (MCSZ) (31–33).

PNKP is a DNA processing enzyme that restores canonical termini at DNA breaks through its 5’ DNA kinase and 3’-phosphatase activities (34–36). Some DNA breaks have noncanonical 3’-phosphate (3’-PO) or 5’-hydroxyl (5’-OH) groups, which cannot be processed by cellular DNA ligases or polymerases. Reactive oxygen species (ROS) produce predominantly DNA breaks harboring 3’-phosphate termini (37), which are also common in DNA breaks induced by ionizing irradiation, stalled topoisomerase I (TOP1) complexes, and as intermediate products of base excision DNA repair (BER). 5’-OH termini are also common products of ionizing radiation or abortive TOP1 reactions (38–42). Although chromatin organization protects DNA from mechanical stresses, its condensation at repairing or replicating DNA sites induces sufficient mechanical stresses to result in DNA breaks (43). Therefore, physical forces are another important mechanism producing DNA breaks which need not have canonical termini. Both 5’-OH and 3’-PO DNA ends are substrates for PNKP.

PNKP deficiencies cause the expected accumulation of DNA breaks, affecting particularly neural progenitor cells, and thus leading to neurodevelopmental and neurodegenerative disorders (44–47). Experimental PNKP depletion causes embryonic death or congenital microcephaly in animal models, depending on the residual levels of enzymatic activities (44, 47). Targeted PNKP deletion restricted to the nervous system resulted in early postnatal lethality in a mouse model. Massive cell death occurred through many still proliferating regions of the developing nervous system. This cell death produced neurodevelopmental defects leading to thinning or even complete loss of the neural cortex (47).

DNA damage is detrimental to genome stability. Therefore, DNA damage signaling activates DNA damage checkpoints to stop cell cycle progression and allow for DNA repair before proceeding into genome replication (DNA synthesis, S phase) or cell division (mitosis, M phase). Cell-cycle progression is driven by the periodical activation and inactivation of the cyclin-dependent kinases (CDK). Activation of CDK1 determines the commitment of a cell to mitosis, whereas its inactivation promotes mitotic exit.

Scheduled activation and inactivation of CDK1 are thus essential for successful mitosis and progression through the cell cycle (comprehensively reviewed in (48)). CDK activity is modulated by regulatory subunits, the cyclins, post-translational modifications, association with other proteins, and subcellular localization. In brief, CDK activation requires binding to their regulatory cyclins, phosphorylation at the activation loop, removal of other inhibitory phosphates, and no inhibition by the CIP/KIP or INK4 CDK inhibitors (48).

Unscheduled mitotic entry in the presence of DNA damage is rare, and it occurs primarily in response to chemo- or radiotherapy in cancer cells defective in checkpoint signaling. Under these conditions, cells undergo catastrophic mitoses producing abnormal nuclear morphologies due to defective chromosome segregation (comprehensively reviewed in (49)).

To prevent mitotic catastrophe, mutagenesis, and other cytopathologies, mitotic entry is highly regulated. It is initiated by the threshold activation of CDK1 (recently reviewed in (50)), which is primarily limited by cyclin availability. CDK1 associates with two types of mitotic cyclins, A or B, which are expressed and degraded in a highly regulated fashion. Cyclin A (CycA) starts to accumulate in S phase and first associates with CDK2, with CycA/CDK activity starting to be detected in early S phase (51). Cyclin B (CycB) starts to accumulate in G2, with CycB/CDK1 activity first detected at mitosis. Both CycA and CycB are degraded in mitosis by regulated ubiquitination-proteasome system. CycA levels decline rapidly after the nuclear envelope breakdown, with degradation completed at prometaphase (52, 53). CycB degradation is initiated by the activated anaphase promoting complex/cyclosome (APC/C) at metaphase. Both CycA and B are thus degraded at the onset of anaphase, rendering CDK1 inactive (reviewed in (54, 55)), and this inactivation is required for mitotic exit. The activity of Cyclin/CDK1 is additionally controlled by inhibitory CDK1 phosphorylations at Thr14 (56–58) and Tyr15 (56, 59) and the activating phosphorylation at Thr161 (60, 61). Phosphorylation on Tyr15, in particular, prevents unscheduled mitotic entry by keeping CycA or B/CDK1 complexes inactive. The CDC25 family phosphatases activate CycB/CDK1 at the onset of mitosis by removing these inhibitory phosphate groups (62–64). Activation of the DNA damage checkpoints inactivates CDC25 phosphatases, thus preventing CDK1 activation and delaying mitotic entry until the DNA damage signaling is turned off.

Both ZIKV and PNKP deficiencies specifically affect human neural progenitor cells, leading to their massive loss during fetal development. Both induce DNA damage and produce chromosomal abnormalities leading to abnormal mitoses. We thus explored whether PNKP could be a cellular factor for ZIKV replication in human iPSC-derived neural progenitor cells (hiNPC) and other cells. We explored potential molecular mechanisms for the direct cytotoxicity of ZIKV to human neural progenitor cells by testing the effects of PNKP inhibition on ZIKV replication and the influence of ZIKV infection on PNKP, the DNA damage operated cell cycle checkpoints, and mitotic entry.

## Materials and methods

### Cell lines

African green monkey Vero E76, human U-2 OS osteosarcoma and HEK-293 embryonic kidney cells were obtained from American Type Culture Collection (CCL-81, HTB-96, CRL-1573, respectively). The CRISPR knockout HCT 116 PNKP-/- human colorectal carcinoma cells (HCT 116 PNKP-/-) were generated and validated before (65). Cells were cultured in DMEM supplemented with 5% FBS at 37°C in 5% CO_2_. Human induced pluripotent stem cell-derived neural progenitor cells (HIP Neural Stem Cells, BC1 line, referred to as ‘hiNPC’) were purchased form MTI-GlobalStem, Inc. (GSC-4311). Cells were grown as monolayers on geltrex-coated surfaces (Geltrex™ LDEV-Free reduced growth factor basement membrane matrix, Gibco, A1413201) in NPC expansion medium: KnockOut™ DMEM/F-12 (Gibco, 12660012) supplemented with StemPro™ neural supplement (Gibco, A1050801), 2 mM (1X) Glutagro (L-Alanine/L-Glutamine dipeptide, Corning), 1X MEM non-essential amino acids (Corning), and 20 ng/mL human heat stable sbFGF (Gibco, PHG0367) and split using accutase (Millipore Sigma, A6964).

### Compounds and treatments

The PNKP inhibitors A12B4C3 and A83B4C63 were synthesized as previously reported (66–68) and kindly provided by Dr. Dennis Hall (University of Alberta). DMSO stocks of A12B4C3 and A83B4C63 (50 mM) were prepared in dimethyl sulfoxide (DMSO), aliquoted and stored at −20°C. Frozen aliquots were thawed fast at 37°C and diluted to target concentrations (30-0.01 μM) in tissue culture media. Roscovitine (Rosco, LC Laboratories, R-1234) stock solution was prepared at 100 mM in DMSO aliquoted and stored at −20°C. Frozen aliquots were thawed fast at 37°C and diluted to 100 μM in pre-warmed DMEM-5% FBS. Compounds were added to cells after removing the inocula and washing with serum free media. Hydrogen peroxide (H_2_O_2_), and ultraviolet irradiation (UV) were used to induce DNA damage. A working solution of 100 μM H_2_O_2_ was prepared by diluting 30% stock solution in serum-free culture media immediately before adding to cells. After 60 min incubation at 37°C in 5% CO_2_, the H_2_O_2_ containing medium was aspirated and cells were washed twice in serum free medium. Complete media (DMEM + 5%FBS or NPC expansion medium) was then added, and cells were returned to the incubator for 1 h. Alternatively cells were washed and overlaid with phenol red free-DMEM, irradiated with UV at 100 J/m^2^ (Spectrolinker XL-1000 UV crosslinker. Spectronics Corp.) and allowed to recover for 1 h in the complete culture medium. Cells were subsequently fixed and processed by immunostaining or lysed and used for Western blotting.

### Viruses

ZIKV strains R103451 (placenta isolate, Human/2015/Honduras) or IbH 30656 (blood isolate, Human/1968/Nigeria) were obtained from BEI resources (NR-50066 and NR-50355, respectively). Stocks were prepared by inoculation of sub-confluent monolayers of U-2 OS cells seeded in a T150 flask 24 h earlier at an MOI of 0.001 in 2 ml DMEM for 1 h. Inoculum was removed and cells were washed twice with 12 ml cold DMEM before overlaying with 15 ml DMEM-5% FBS. Infected cells were incubated for 24 h at 37°C in 5% CO_2_ to reach confluence, and then split 1:3. Supernatants were harvested 5 days post inoculation and cells were overlaid with 15 ml DMEM-5%FBS. Typically, 3-4 harvests were collected every 4-5 days before cells reached full cytopathic effect. Collected medium was clarified by centrifugation at 3,000 g for 10 minutes at 4°C. Cleared culture media was concentrated 10 times for each harvest, using 50 mL 100K Amicon Ultra centrifugation tubes spun at 3500 g for 25 minutes at 4°C. Retentates were aliquoted into 500 μL, frozen in ethanol-dry ice bath and stored at −80°C.

### Antibodies

Primary and secondary antibodies are listed in Supplementary Tables 1 and 2 (**Suppl. Tab.1, 2**)

### ZIKV infections

For immunofluorescence (IF), 3×10^5^ (hiNPC) or 1.5×10^5^ (U-2 OS, Vero, 293) cells were seeded onto geltrex-coated (hiNPC) or uncoated (U-2 OS, Vero, HEK-293) 21 mm diameter round coverslips in 12-well plates and infected 24 h later with 1-3 foci forming units (FFU) (hiNPC) or 1-3 plaque forming units (PFU) (other cells) per cell of ZIKV in 150 ml of expansion medium (hiNPC) or DMEM (other cells). Inoculated cells were incubated for 1 h at 37°C, rocking and rotating every 10-15 min. Cells were washed once with 2 ml/well cold KnockOut DMEM:F12 (hiNPC) or DMEM (other cells) before overlaying with 1.5 ml/well 37°C complete expansion medium (hiNPC) or DMEM-5% FBS (other cells) and incubated at 37°C in 5% CO_2_. Cells were fixed and processed by immunostaining at pre-selected times.

For drug treatments, cells were seeded on geltrex-coated 12-well plates as described and infected 24 h later with 3 FFU (hiNPC) or 3 PFU (other cells) per cell of ZIKV as described. Culture media were harvested at pre-selected times, cleared by centrifugation at 3,000 g for 10 minutes at 4°C, and titrated immediately or frozen at −80°C for later titration.

For Western blotting or immuno-kinase activity assays, 3×10^6^ cells were seeded on T75 flasks or 10 cm diameter dishes and infected 24 h later with 3 FFU (hiNPC) or 3 PFU (U-2 OS) per cell of ZIKV in 1 ml serum free medium, as described. Cell monolayers were detached with accutase (hiNPC), or trypsin (U-2 OS) at pre-selected times, pelleted, resuspended in 1 ml ice-cold PBS (10 μl aliquots were used for cell counting) and pelleted again. PBS was aspirated and cell pellets were frozen at −80°C until further processing.

### Plaque forming assays

Vero cells (1.2x 10^5^) were seeded into each well of 12-well plates and incubated at 37°C in 5% CO_2_. Cells were inoculated with 150 μl of sample the following day and incubated for 1 h at 37°C, rocking and rotating every 10 min. Cells were washed twice with 2 ml/well cold DMEM before overlaying with 1.5 ml/well 37°C DMEM-5% FBS containing 1% methylcellulose and incubating at 37°C in 5% CO_2_ until plaques developed, typically 4-6 days. Cells were fixed and stained overnight with 0.5% crystal violet in 17% methanol. Plaque numbers are expressed as PFU/ml for titration of viral stocks or normalized to infected cell number and expressed as PFU/cell (virus burst) for dose response or time of addition analyses.

### Foci formation assays

ZIKV infectious titers in hiNPC we evaluated by FFU. hiNPC (3.0 x 10^5^ cells/well) were seeded into Geltrex-coated 12-well plates in 1 ml of NPC expansion medium and incubated at 37°C in 5% CO2. Cells were inoculated 24 h later with 150 μl per well of serial dilutions of ZIKV stocks in the expansion medium and incubated at 37°C for 1 h, rocking and rotating every 10-15 min. Inocula were removed and cells were washed twice with 1.5 ml/well cold unsupplemented KnockOut DMEM:F12. Cells were overlaid with 1 ml/well expansion medium, containing 0.75% methylcellulose and incubated for 48 h at 37°C and 5% CO_2_. Cells were fixed and processed for indirect immunofluorescence for flavivirus group E antigen (clone 4G2). Calculated foci numbers were expressed as FFU/ml.

### Time of addition assay

hiNPC cells were infected with 3 FFU of ZIKV/cell. After 1 h adsorption, cells were washed and overlaid with 2 ml drug-free expansion medium or expansion medium containing 100 μM Rosco. Drug-free expansion medium was removed at 3, 6, 9, 12, or 15 hours post infection (hpi) from replicate series of infected monolayers and replaced with 2 ml of expansion medium containing 100 μM Rosco. An additional series of infected monolayers was incubated continuously in vehicle (0.1% DMSO) containing drug-free expansion medium. For each time of addition, culture media were sampled just before Rosco addition and at the end of experiment (24 hpi), to calculate viral burst from the time of drug addition. To this end, 100 μl of medium was removed and substituted with 100 μl fresh expansion media. Samples were frozen at −80°C until titrated on Vero cells. For untreated infections, culture medium was equally sampled at every time-point from 1 to 15 h, and then at 24 h. Viral burst, as percent of burst in vehicle control, is plotted against hours post infection (hpi). Each time point indicates the average and range of two independent experiments.

### Immuno complex kinase assay

Cell lysates were prepared by resuspending cell pellets to 1×10^4^ cells/μl in non-denaturing lysis buffer (50 mM Tris-HCl, pH 7.4, 1 mM EDTA, 1 mM EGTA, 150 mM NaCl, 1% Triton X-100), supplemented with protease and phosphatase inhibitor cocktails (cOmplete, EDTA-free and PhosSTOP from Roche) and incubated for 30 min at 4°C. Lysates were cleared by centrifugation (15,000 g, 15 min, 4°C) before 100 μl of supernatant, equivalent to 1×10^6^ cells, was incubated with 25 μl of agarose-coupled anti-CDK1 or anti-CycA2 antibodies (sc-54AC or sc-239AC, Santa Cruz Biotechnology, respectively) for 2 h at 4 °C with gentle rocking. After three washes in lysis buffer and one wash in kinase buffer (50 mM Tris-HCl, pH 7.4, 10 mM MgCl_2_, 1 mM DTT), kinase assays were performed by incubating the immunoprecipitated proteins and beads in 30 μl kinase buffer supplemented with 50 μM ATP and 3 μg histone H1 (EMD Millipore, 14-155) for 30 min at room temperature. Kinase activity was measured using ADP-Glo Kinase assay (V6930, Promega) according to the manufacturer’s instructions. Luminescence was measured using a TECAN infinite 200Pro plate reader. Kinase activity, proportional to relative luminesce units (RLU), was plotted against time after infection.

### Indirect immunofluorescence

For confocal imaging, cells were fixed with 3.7% buffered formaldehyde for 15 min at room temperature. Fixative was removed, cells were washed three times with 1 ml PBS each, permeabilized with 0.5% Triton X100 in PBS for 15 minutes at RT, and blocked with 500 μl 5% normal goat serum (NGS) in PBS for 1 h at room temperature (RT). Primary antibodies (**Suppl. Tab. 1**) were diluted in 1% NGS in PBS and incubated overnight at 4°C. Primary antibodies were removed and cells were washed three times in 1 ml of 0.2% TritonX100 in PBS each time. Fluorescently-conjugated secondary antibodies (**Suppl. Tab. 2**) were diluted 1:1,000 in 1% NGS in PBS, incubated for 1 h at room temperature protected from light and washed 3 times in 1 ml each wash of 0.2% TritonX100 in PBS.

Cells were counterstained with 2 μg/ml DAPI in PBS for 5 min at room temperature. Coverslips were removed from the wells, rinsed briefly in dH_2_O, and mounted onto glass microscope slides using 10 μl Prolong Gold (Molecular Probes, P10144) or Prolong Glass (Invitrogen, P36984) antifade mounting media. The slides were protected from light until imaging by confocal microscopy using a Zeiss LSM 510 or an Olympus FV 3000 microscope and ZEN2.3ProHWL or FV315-SW image acquisition software, respectively. ImageJ software (69) was used to convert raw image stacks to single channel (grey scale) or merged (color scale) images and to add scale bars.

### Metabolic labeling with EdU

hiNPC (2.5 x 10^5^) were seeded on Geltrex-coated 21 mm diameter glass coverslips in 1.0 ml growth medium. 24 hours after plating, 1.0 ml of 20 μM EdU in growth medium was added (10 μM final concentration) and cells were returned to the incubator for a 15 min pulse labelling. Cells were subsequently fixed and permeabilized for IF as described above, blocked with 1 ml of 3% BSA in PBS, washed and stained using Click-iT™ EdU Cell Proliferation Kit for Imaging, using Alexa Fluor™ 488 dye (Thermo Fisher Scientific, C10337), according to the manufacturer’s protocol. Cells were subsequently stained for ZIKV NS1 (IF) and counterstained with DAPI.

### Cytotoxicity

Cytotoxicity and cytostasis in hiNPC cells were tested using CellTiter-Glo® luminescent cell viability assay (Promega, G7570). Cells (5 x 10^3^ cells/well) were seeded in 96-well plates to reach 25% confluence at 24 h after plating. Test compound dilutions were added 24 hours later in 150 μl in hiNPC maintenance medium and incubated for 2 to 3 days at 37°C. Relative cell number was determined daily by incubating replicated monolayers with CellTiter-Glo reagent diluted in phenol-red free DMEM for 30 min at RT. Luminescence was measured using TECAN infinite 200Pro plate reader. Relative cell number was plotted against time of treatment for each drug dilution. Cytostasis was evaluated by no increase or decrease in the number of viable cells during the incubation period.

### Western blotting

Cell monolayers in T75 flasks or 10 cm diameter dishes were detached with accutase (hiNPC) or trypsin (U-2 OS), pelleted, and resuspended in 1 ml of ice-cold PBS (10 μl samples were collected for counting cells) and pelleted again. PBS was aspirated and cell pellets were frozen at −80°C. Sufficient ice-cold lysis buffer (50 mM Tris-HCl, pH 7.4, 1 mM EDTA, 1 mM EGTA, 150 mM NaCl, 1% Triton X-100, 0.5% sodium deoxycholate, 0.5% SDS) supplemented with protease and phosphatase inhibitor cocktails (cOmplete, EDTA-free and PhosSTOP from Roche) was added to the pellets to yield 1×10^4^ cell equivalent/μl. Samples were incubated for 30 min at 4°C with rocking and then sonicated (3 x 30 sec interval, high setting in a Bioruptor water bath sonicator, Diagenode). Lysates were cleared by centrifugation at 13,000 g for 15 min at 4°C. Supernatants were collected, mixed 4:1 with reducing NuPAGE™ LDS sample buffer 4x (ThermoFisher Scientific, NP0007) and boiled for 10 minutes. Proteins were separated by electrophoresis in 12 or 7% polyacrylamide gels (the latter for high molecular weight proteins, ATM and ATR) and transferred to polyvinylidene fluoride or nitrocellulose membranes. Membranes were blocked with TBS containing 0.1% Tween-20 (TBST) supplemented with 5% BSA. Primary antibodies (**Suppl. Tab. 1**) diluted in TBST were added and incubated overnight at 4°C with rocking. Membranes were washed with TBST three times for 10 minutes each and incubated with the corresponding fluorescently-labeled secondary antibody (**Suppl. Tab. 2**) for 1 h at room temperature with gentle rocking. Membranes were washed as described and signal was detected and recorded using a ChemiDoc MP imager (BioRad).

## Results

### Two PNKP inhibitors, A83B4C463 and A12B4C3, inhibit ZIKV replication, as does knocking out PNKP

PNKP is an essential DNA damage repair enzyme which, like ZIKV infection, is also linked to microcephaly (31, 32). We asked whether ZIKV infection may affect, or be affected by, PNKP. To this end, we tested the effect of two PNKP phosphatase inhibitors, A83B4C63 and A12B4C3, on ZIKV replication in hiNPC cells. We treated already infected cells with semi-logarithmic dilutions of the inhibitors and titrated the cell-free infectivity at 24 h. Both inhibitors induced dose response inhibition of ZIKV replication (Fig. 1A). A83B4C63 and A12B4C3 were no cytotoxic or cytostatic at up to 30 μM or 10 μM, respectively, in that cells continuously doubled for at least two days, whereas A12B4C3 was cytostatic (but not cytotoxic) at 30 µM (Fig. 1C). Inhibition of PNKP phosphatase activity in cells with concentrations of A12B4C3 and A83B4C63 in the same range of 1 to 10 µM inhibit the repair of single and double stranded DNA breaks and sensitizes cancer cells to the topoisomerase poison camptothecin and γ-irradiation (66-68, 70-72).

**Figure 1.**
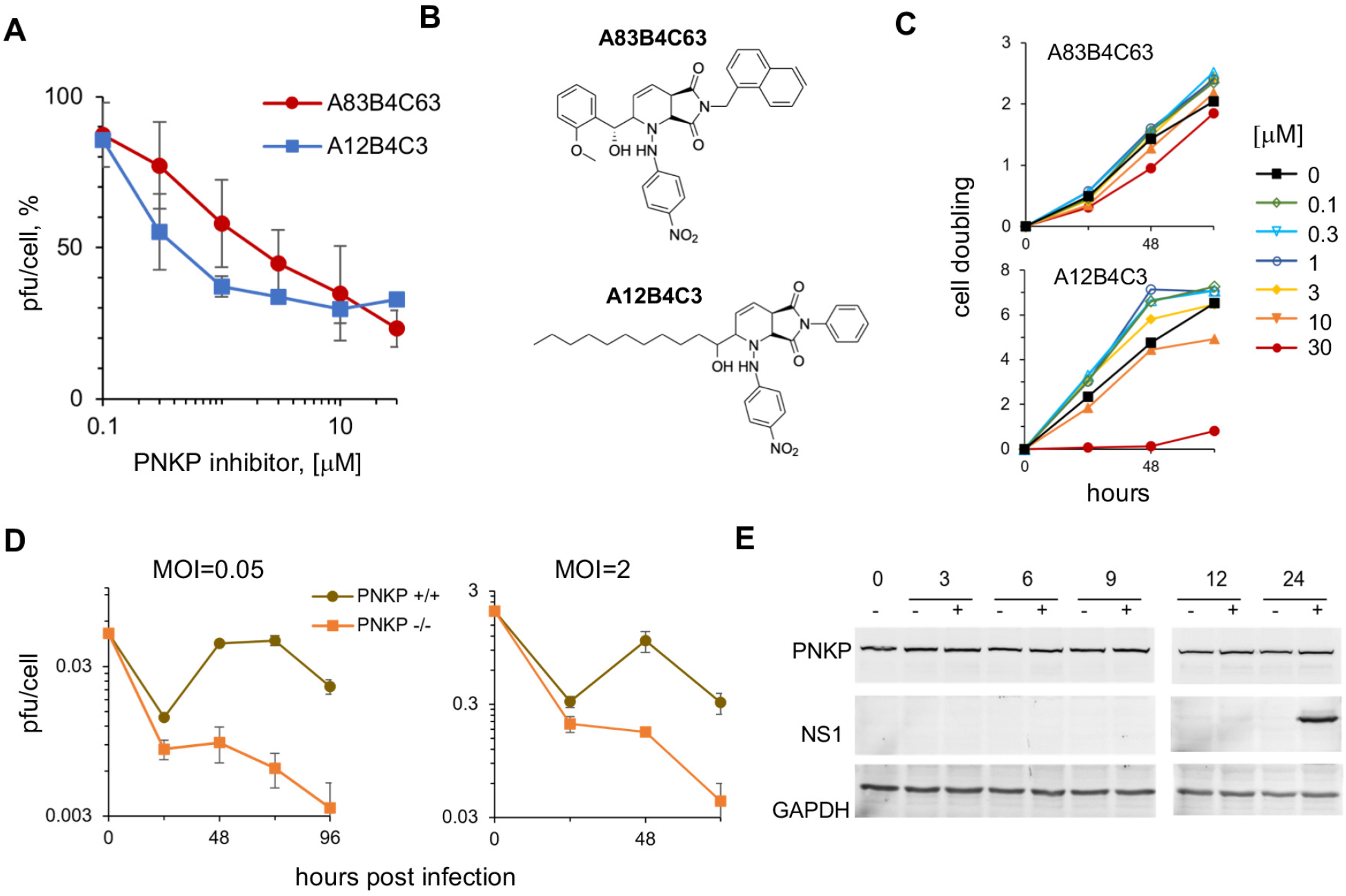
Two PNKP phosphatase inhibitors or PNKP genetic knockout inhibit ZIKV replication. (A) hiNPC were inoculated with ZIKV (MOI=0.1), washed, and overlaid with media containing the PNKP inhibitors A83B4C3 or A12B4C3, or DMSO. Culture supernatants at 48 hpi were titrated; results expressed as percent PFU/cell relative to DMSO-treated infections; n=2; (average +/- range). (B) Chemical structures of the PNKP phosphatase inhibitors A83B4C63 and A12B4C3. (C) hiNPC were seeded in 96-well plates to reach 25% confluence at 24 h after plating, when test dilutions of the PNKP inhibitors were added (T_0_). Relative cell numbers were evaluated daily. Results show cell doublings relative to T_0_, n=1. (D) WT or PNKP knockout HCT 116 cells were inoculated with ZIKV (MOI=0.05 or 2), washed, and overlaid with media. Culture supernatants were collected every 24 h and titrated on Vero cells; results expressed as PFU/cell; n=2; (average +/- range). (E) hiNPC infected with ZIKV R1034 (MOI=3) or mock infected were harvested at the indicated times after infection, lysed, and analyzed by WB (whole cell lysates). Immunoreactive bands were visualized by fluorescently labelled secondary antibodies.

Based on these results we proposed that PNKP phosphatase activity was required for ZIKV replication. We tested this model by evaluating ZIKV replication in a CRISPR knockout HCT 116 PNKP-/- cell line (65). We infected WT and PNKP-/- HCT 116 cells with ZIKV at low or high multiplicity of infection (MOI=0.05 or 2, respectively). We harvested culture supernatants every 24 h for up to 120 h and titrated the cell free infectivity. ZIKV failed to replicate in the PNKP knockout cells infected at either low or high multiplicity. The ZIKV replication cycle had been completed in 72 or 48 hours in the WT cells infected at low or high MOI, respectively (Fig. 1D), and the viral titers decreased thereafter.

PNKP is constitutively expressed, and its levels are regulated by proteasomal degradation (73). Host factors required for viral replication are often upregulated during infection. However, PNKP levels were constant during ZIKV infection and did not differ from those in mock-infected cells (Fig. 1E).

### PNKP relocalizes to the cytoplasm, where it co-localizes with NS1 during ZIKV infection

PNKP normally localizes to the nucleus and mitochondria (74) whereas ZIKV replicates in cytoplasmic ER-derived membranous structures (26). We thus evaluated PNKP subcellular localization in ZIKV-infected cells (Fig. 2A). PNKP localized predominantly to the nucleus in mock infected hiNPC, showing a disperse staining pattern, as expected (74). In contrast, PNKP signal was concentrated in the cytoplasm at the center of ZIKV-infected cells, whereas abnormal and often multi-lobular nuclei occupied the cell periphery. Cytoplasmic PNKP colocalized with NS1 (PCC, 0.83-0.88, Fig. 2A). PNKP colocalized in the cytoplasm with NS1 in about 80-95% of cells at 24 hpi. PNKP signal started to be depleted from the nucleus at 3 h post infection, with a gradual increase in cytoplasmic staining observed from 6 hpi. PNKP colocalization with NS1 was obvious as early as ZIKV replication factories were detected at 12 hpi, although it was detectable as soon as 9 hpi in some infected cells staining positive for NS1 (Fig. 2C).

**Figure 2.**
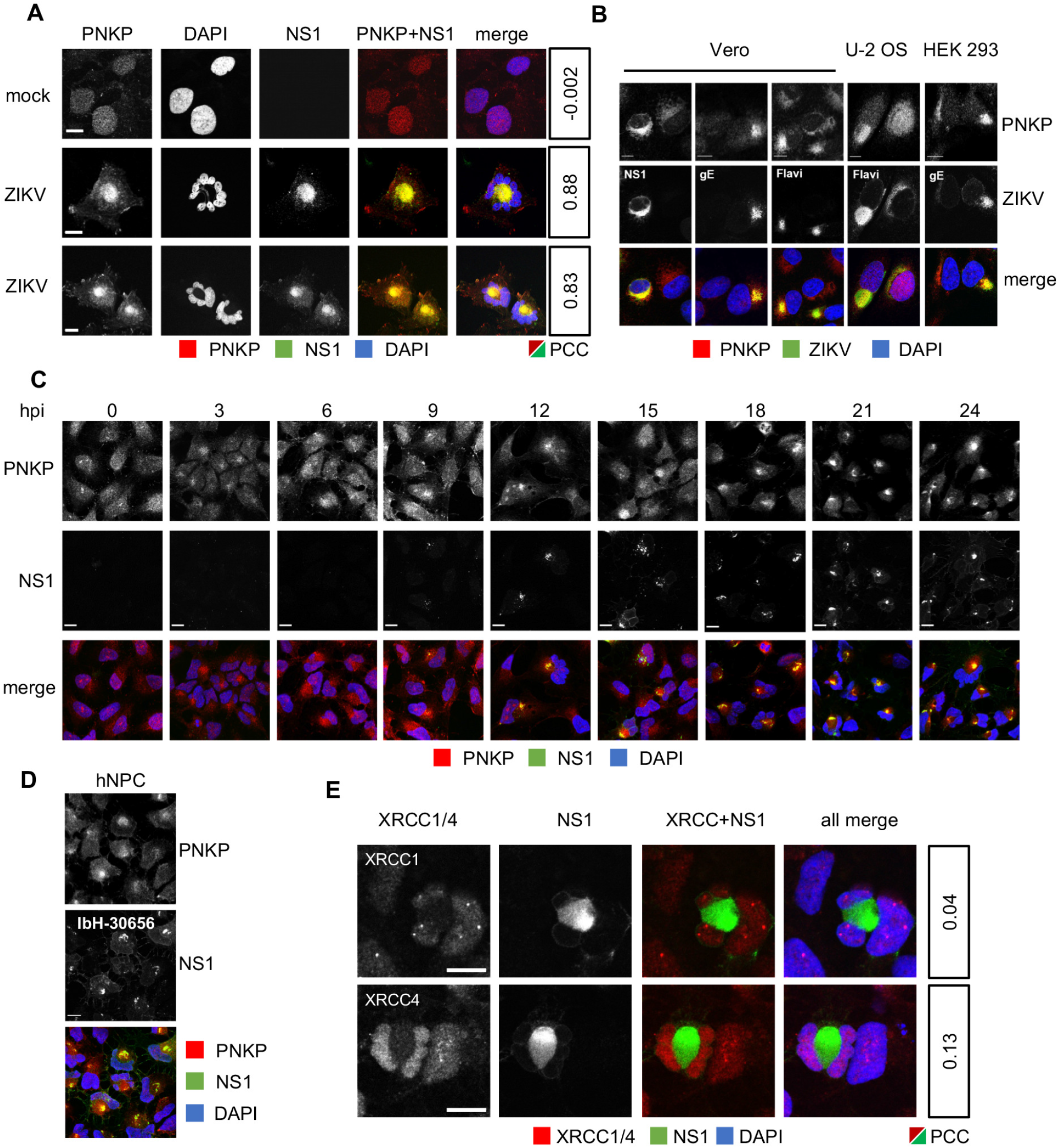
PNKP relocalizes to the cytoplasm in ZIKV-infected NPC, where it colocalizes with NS1. (A, B, E) hiNPC, Vero, U-2 OS or HEK 293 cells were infected with ZIKV R1034 (MOI=1), fixed 24 h later, and analyzed by IF for NS1 and PNKP (A, B) or XRCC1 or XRCC4 (E). Pearson’s colocalization coefficients between NS1 and PNKP or XRCC1 and XRCC4 signals were calculated using the colocalization finder plugin in ImageJ. (C) hiNPC infected with ZIKV R1034 (MOI=2) were fixed at the indicated times after infection and analyzed by IF for NS1 and PNKP. (D) hiNPC were infected with ZIKV IbH-30656 (MOI=1), and 24 h later fixed and analyzed for PNKP and NS1 (IF). Images representative of at least 2 independent experiments; scale bars, 10μm.

Co-localization of PNKP and NS1 was not unique to neural progenitor cells. Cytoplasmic PNKP accumulation was also detected in three non-neural cell lines permissive to ZIKV replication, U-2 OS, Vero, and HEK293 (Fig. 2B). Notably, however, the nuclear morphology in these non-neural progenitor cells was much closer to that of their non-infected counterparts. PNKP localization to ZIKV RFs was detected using three different ZIKV-specific antibodies recognizing ZIKV NS1 or gE, or flavivirus group E antigen (Fig. 2B).

PNKP relocalization was not unique to the contemporary ZIKV strain R103451 (Honduras/2016/human placenta). Cytoplasmic PNKP and abnormal nuclear morphology were equally induced in neuronal progenitor cells by historical ZIKV strain IbH-30656 (Nigeria/1968) (Fig. 2D).

### ZIKV-induced PNKP relocalization is independent of XRCC1 or XRCC4

PNKP is recruited to nuclear DNA damage sites by interactions with the DDR scaffold proteins XRCC1 or XRCC4 (75–77). Depending on the type of DNA damage, XRCC1 (single strand breaks) or XRCC4 (double strand breaks) organize multiprotein complexes at DNA lesions to concentrate DNA repair enzymes, including PNKP, DNA ligases, and polymerases, along with multiple regulatory and signaling proteins. We thus asked whether PNKP was recruited to the cytoplasm by XRCC1 or XRCC4. XRCC1 and XRCC4 are normally nuclear and both remained nuclear in ZIKV-infected hiNPC. There was no co-localization of either with NS1 (PCC 0.13-0.04) (Fig. 2E). XRCC1 or XRCC4 therefore do not account for the cytoplasmic PNKP accumulation.

### ZIKV infected hiNPC accumulate DNA damage

Downregulation or inhibition of PNKP sensitizes cells to genotoxic stress and results in accumulation of unrepaired DNA (45-47, 66, 67, 70, 71). Cytoplasmic PNKP sequestration in ZIKV infection may result in depletion of functional nuclear PNKP, with the consequent accumulation of DNA damage. We used γH2AX staining to test for DNA damage. H_2_O_2_ treatment of uninfected cells induced the expected canonical, bright, well defined, and isolated nuclear γH2AX foci, detectable at 10 min post treatment (Fig. 3A, 3C and **suppl. Fig. 1A, 1B**). Infected cell nuclei stained positive for γH2AX at 24 hours after infection (Fig. 3A). Some baseline level of γH2AX staining was detected also in mock-infected cells or at time zero after ZIKV infection, but the nuclear γH2AX foci started to obviously increase in number, size, and brightness above baseline after infection (**Suppl. Fig. 1A**), and most ZIKV-infected cells had strong nuclear γH2AX signal at 32 and 48 h. Thus, ZIKV infection induces DNA damage that continues to accumulate during infection.

**Figure 3.**
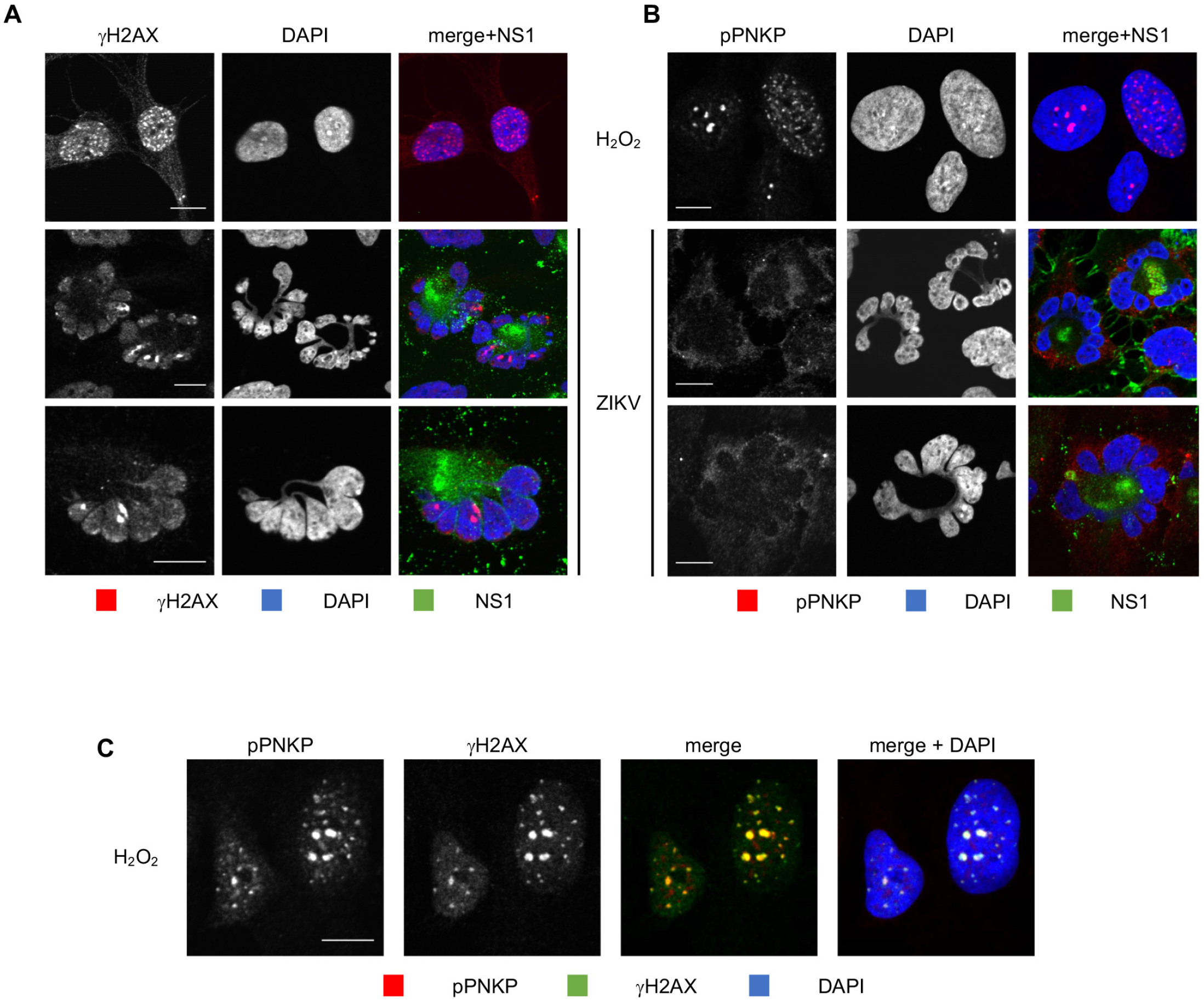
ZIKV infected NPC accumulate DNA damage but fail to accumulate phosphorylated PNKP. (A, B) hiNPC infected with ZIKV (MOI=1) were fixed at 26 hpi and analyzed by IF for NS1 and γH2AX (A) or pPNKP (B). (C) Mock-infected cells were treated with 100 μM H_2_O_2_ for 1 h, washed, overlaid with fresh complete medium, fixed 1 h later, and analyzed by IF for γH2AX and pPNKP; scale bars, 10 µm; images representative of 3 independent experiments.

### PNKP is not phosphorylated at the DNA damage sites in ZIKV-infected cells

In response to DNA damage signaling, PNKP is recruited to the DNA breaks where it is phosphorylated by ATM. Phosphorylation allows for the accumulation of PNKP at the DNA breaks, accumulation which is necessary for efficient repair (73, 78, 79). ZIKV-induced accumulation of DNA damage could be the consequence of nonfunctional repair mechanisms due to the cytoplasmic PNKP localization (Fig. 2A). However, cytoplasmic PNKP could still be recruited to the nucleus, or a small population of PNKP may remain in the nucleus and be activated. We tested PNKP phosphorylation using phospho-specific PNKP antibodies (Ser114). pPNKP accumulated in scattered nuclear foci in uninfected cells treated with H_2_O_2_ (Fig. 3B, C), as expected. Also as expected, H_2_O_2_-induced PNKP foci were detected at 30 min post treatment and colocalized with γH2AX foci (Fig. 3C and **suppl. Fig. 1B**). In contrast, there was no detectable p-PNKP signal in ZIKV-infected cells (Fig. 3B, **Suppl. Fig. 3C**). No pPNKP signal was detected at any time during ZIKV infection **(Suppl. Fig. 3C).**

### ZIKV infection of hiNPC induces obvious gross mitotic abnormalities

Throughout all experiments (Fig. 2A, B, C; 3A, B; **Suppl. Fig. 1A, C**), we had noted that ZIKV infection consistently induced multilobed nuclei, micronuclei, and chromosomal bridges in hiNPC but not in other cells (Fig. 2D, 4A). These defects are all consistent with mitotic abnormalities. We analyzed the relative frequency of the putative mitotic aberrations in hiNPC or U-2 OS. Grossly abnormal nuclear morphologies were more frequent, and more severe, in infected hiNPC than U-2 OS (Fig. 4A, B), Vero, or HEK293 cells (Fig. 2D). These abnormal nuclei did not show the typical features of apoptosis; they rather resembled the nuclear morphology of cells undergoing mitotic catastrophe (MC) (80, 81). The morphological hallmarks of MC are micro and multi-nucleation with the nuclear envelope assembled around individual clusters of mis-segregated chromosomes (81, 82). Micro and multi-nucleation were both common features among ZIKV-infected hiNPC (Fig. 4D), and lamin B1 staining revealed that the individual nuclear lobules and micronuclei were indeed surrounded by nuclear envelopes (Fig. 4D). ZIKV induced obvious gross nuclear abnormalities in hiNPC from 12 hpi (Fig. 4E), and the nuclei of infected hiNPC were globally and grossly abnormal, and DNA was largely no longer replicating, at 24 hpi (Fig. 4F).

**Figure 4.**
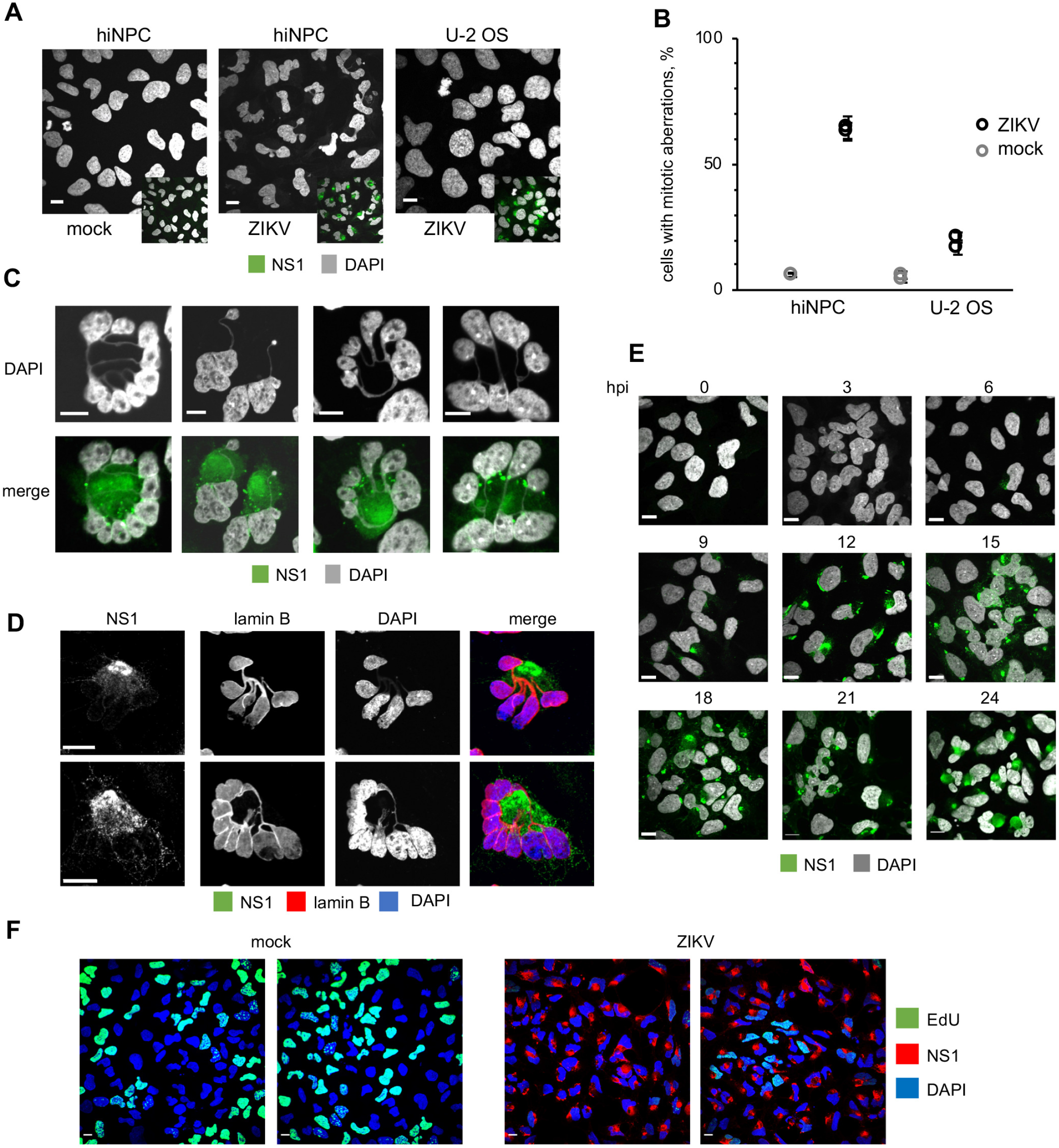
ZIKV induces grossly abnormal nuclear morphologies in hiNPC, morphologies which are consistent with the hallmarks of mitotic catastrophe. (A) hiNPC or U-2 OS were infected with ZIKV R1034, (MOI=1), fixed at 24 hpi and analyzed by IF for ZIKV NS1. (B) Cells with obvious and gross nuclear aberrations at 24 h were counted in 5 wide field confocal images. Results expressed as percentage of all cells (for mock infections) or infected cells (for ZIKV infections); n=2 (average, +/- SD for each individual experiment). (C) Infected hiNPC, as identified by NS1 staining at 24 hpi and counterstained with DAPI were analyzed using high magnification confocal imaging. (D) hiNPC were infected with ZIKV R1034, (MOI=1), fixed at 24 hpi and analyzed by IF for NS1 and lamin B. (E) hiNPC were infected with ZIKV R1034, (MOI=3), fixed at indicated times, and analyzed by DAPI staining and IF for NS1. (F) hiNPC were infected with ZIKV R1034, (MOI=3), or mock-infected, for 24 h were metabolically labelled for 15 minutes with EdU, fixed, and stained for EdU and NS1 (IF). Scale bars, 10 μm; images representative of 2-4 independent experiments.

### ZIKV infected human neural progenitors undergo mitotic catastrophe

MC results from unscheduled mitotic entry in the presence of DNA damage or unfinished DNA replication with the consequent incomplete decatenation. As a result, chromosomes cannot be properly segregated, and pulling by the mitotic spindle results in DNA damage and chromosomal bridges, and consequent mis-segregation of chromosomes. The end results are abnormal multi-lobular nuclei. As a consequence of the resulting gross genomic defects, cells die at mitosis or at subsequent cell cycles. If ZIKV-induced nuclear abnormalities are indicative of MC, then they should be prevented by blocking mitosis. To test for this possibility, we used roscovitine (Rosco), which inhibits mitosis by inhibiting CDK1, and also CDK2 (it also inhibits other protein kinases not involved in mitosis like CDK5) (83, 84). At 100 μM (85–87), Rosco inhibited hiNPC cell doubling, which results from inhibition of mitosis, without decreasing viability (Fig. 5A). ZIKV-infected cells treated with DMSO vehicle showed morphological features characteristic of MC, as previously observed (Fig. 2A, Fig. 3A, B, Fig. 4C, D), whereas Rosco-treated infected cells, did not (Fig. 5B). The nuclear morphology of infected cells treated with Rosco was closer to that of mock infected cells than to that of the untreated infected ones, with far fewer cells presenting multilobular nuclei (MLN) or chromosomal bridges (CHB) (Fig. 5C). The frequency of MLN in infected cells decreased from about 60-70% to 10-15%, and that of CHB from 35-45% to about 5%. Mock-infected cells had less than 10% MLN or less than 1% CHB regardless of treatment. Inhibiting cell duplication with an inhibitor of CDK1 (and also of other protein kinases like CDK2, CDK5 and CDK7) thus prevents the abnormal nuclear morphology induced by ZIKV. Unexpectedly, no ZIKV RF were obvious in Rosco-treated cells, as evidenced by the lack of clusters of cytoplasmic NS1 signal (Fig. 5B, lower panel). NS1 accumulation was delayed by some 6 hours by Rosco, in that there was about as much NS1 at 24 hpi in the treated cells as at 18 hpi in the untreated ones (Fig. 5D).

**Figure 5.**
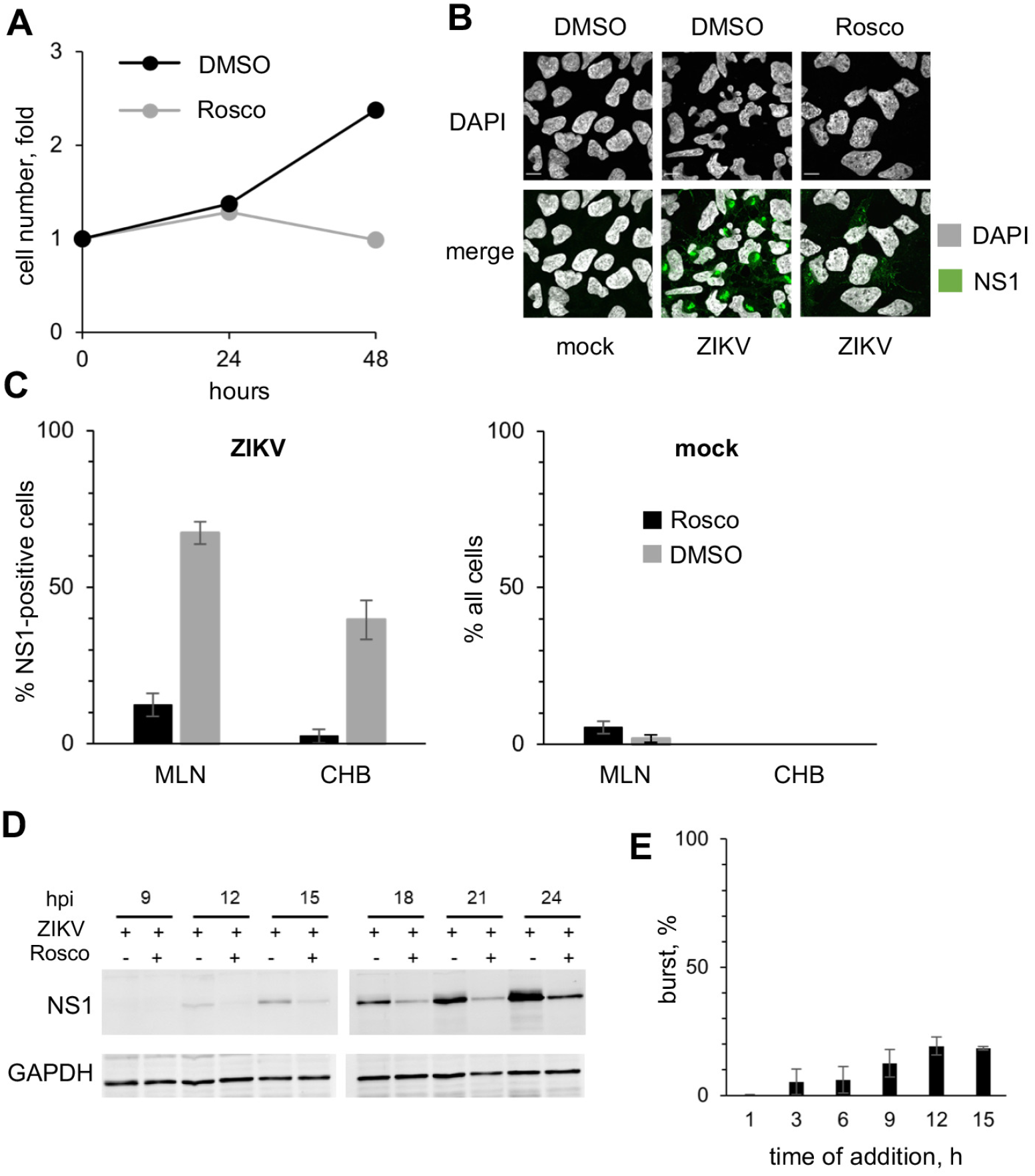
Inhibition of cell doubling with roscovitine inhibits the aberrant nuclear morphologies and ZIKV replication. (A) hiNPC were seeded in 96-well plates to reach 25% confluence at 24 h after plating, when 100 µM Rosco or vehicle (DMSO) were added (T_0_). Relative cell numbers were evaluated daily. Results show cell doublings relative to T_0_, n=1. (B-E) hiNPC were infected with ZIKV R1034 (MOI=3), or mock-infected, washed, and overlaid with media containing 100 µM Rosco or vehicle (DMSO) at the time of inoculum removal (B, C, D) or at indicated time post infection (E). (B) Cells were fixed at 24 hpi and analyzed by IF; images are representative of 2 independent experiments. (C) ZIKV or mock-infected cells with multilobular (≥3 lobules) nuclei (MLN) or chromosomal bridges (CHB), counted in the wide field confocal images from (B); results expressed as percent of NS1-expressing (ZIKV-infected) or all (mock-infected) cells; n=6 (average +/- range). (D) Cells were lysed at indicated time points and analyzed by Western blot; images are representative of 3 independent experiments. (E) ZIKV-infected cells were treated with Rosco at the indicated times after infection. Culture supernatants at time of drug addition and at 24 hpi were titrated on Vero cells; results expressed as burst from time of treatment relative to vehicle-treated infections; n=2 (average +/- range).

### Roscovitine inhibits ZIKV replication

Given the lack of obvious NS1 positive ZIKV RF (Fig. 5B) and delayed NS1 accumulation (Fig. 5D) in Rosco-treated infected cells, we tested the effect of Rosco added at selected times after infection on ZIKV replication.

Cells were infected with 3 PFU of ZIKV/cell. After 1 h adsorption, the inoculum was removed and monolayers were washed and overlaid with drug-free medium or medium containing 100 μM Rosco. The drug-free medium from replicate infections was replaced every 3 h with medium containing 0 or 100 μM Rosco. The infectivity released to the culture medium by Rosco- or vehicle-treated cells was sampled every 3 h from the time of compound or vehicle addition until 24 hpi. The results are expressed as virus burst from the time of Rosco addition to 24 hpi and presented as percentage of vehicle-treated controls. Addition of Rosco immediately after adsorption resulted in about 99.7% inhibition of ZIKV replication (Fig. 5E). The addition of Rosco at 3-6 h allowed only for low levels of replication, about 5-6% of those in untreated infections. Even when Rosco was added at 9, 12 or 15 hpi, ZIKV replicated to only 12, 19 and 18% of the levels in untreated infections, respectively. Thus, Rosco inhibits ZIKV replication even when added at 15 hours after infection, and with similar potency as it inhibits the nuclear replicating DNA viruses HSV-1, HCMV, EBV, or VZV (85, 88–92).

### ZIKV induces cytoplasmic CDK1 accumulation

Membrane limited cytoplasmic organelles undergo rapid morphological changes in size and shape as cells enter mitosis, fragmenting and dispersing to enable their correct inheritance between the two daughter cells (reviewed in (93)). Following the nuclear pore complex (hiNPC) disassembly at prophase, nuclear envelope membrane proteins are released into the mitotic endoplasmic reticulum (ER), which is remodeled from predominantly sheets in the interphase into predominantly tubules or cisternae in mitosis (94, 95). This entire process is initiated by the cytoplasmic activity of mitotic CDK1.

ZIKV induces a large remodeling of the ER to assemble cytoplasmic RF. ZIKV-remodeled ER membranes consist of mostly tubules and cisternae, globally like the mitotic ER, and Rosco, which inhibits mitosis by mostly inhibiting CDK1, and also CDK2, inhibited the formation of the RF (Fig. 5B). RF assembly may thus require CDK1 activity. We therefore asked whether ZIKV might recruit CDK1 to induce the ER membrane rearrangements required for RF assembly.

CDK1 localization varies during the cell cycle. In uninfected cells, CDK1 was dispersed throughout the cytoplasm and nucleus during interphase and started to accumulate at the nuclear periphery and centrosomes at the beginning of prophase, as expected (96).

Later in mitosis, CDK1 localized around condensed chromosomes (Fig. 6A). In ZIKV-infected cells however, there was a cytoplasmic accumulation of CDK1 (Fig. 6C, upper panel) starting at 3 hours after infection, independent of any mitotic changes in nuclear morphology (Fig. 6E). CDK1 signal was enriched around NS1, but CDK1 and NS1 signals did not fully colocalize (PCC 0.62, Fig. 6C, upper panel, **D, E**) and were rather intermixed, in contrast to the co-localization between PNKP and NS1 (Fig. 6D).

**Figure 6.**
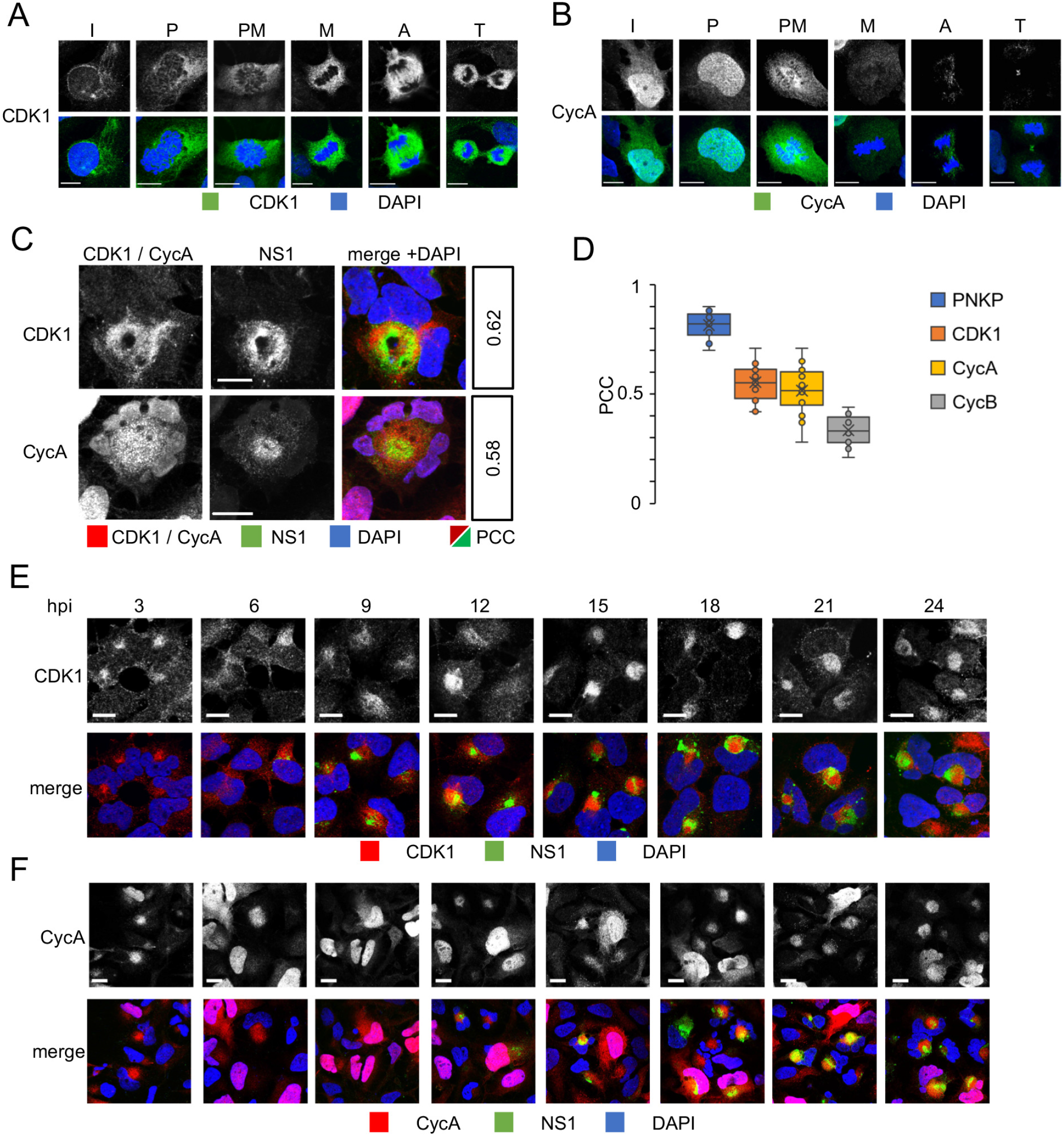
ZIKV infection induces cytoplasmic accumulation of CDK1 and Cyclin A. (A, B) uninfected hiNPC were fixed 24 h post plating and analyzed by DAPI staining and IF for CDK1 (A), or CycA (B). Cells in different cell cycle phases were identified based on nuclear DNA. I, interphase; P, prophase; PM, prometaphase; M, metaphase; A, anaphase; T, telophase. (C-F) hiNPC were infected with ZIKV R1034, (MOI=2), fixed at 24 hpi (C), or at indicated times (E, F) and analyzed by IF for NS1 and CDK1 (C, E) or CycA (C, F). Scale bars, 10 µm. (D) hiNPC were infected with ZIKV R1034, (MOI=2), fixed at 24 hpi and analyzed by IF for NS1 and PNKP, or CDK1, or CycA, or CycB. Pearson’s colocalization coefficients between NS1 and PNKP, CDK1, CycA, or CycB calculated from single cell confocal images (n=18) using the colocalization finder plugin in ImageJ. Results expressed as box plots, cross mark, mean; boxes, 25th to 75th percentile; whiskers, 5th and 95th percentiles.

### ZIKV induces accumulation of cytoplasmic cyclin A

CDK1 activity is highly regulated at multiple levels. Binding of cyclin A or B, cellular localization of the cyclin/CDK1 holoenzymes, several activating and inhibitory phosphorylations, and binding by specific inhibitors, all regulate CDK1 activity to prevent unscheduled mitotic entry and ensure successful exit from mitosis. ZIKV infection induced MC and abnormal cytoplasmic accumulation of CDK1. We thus asked whether either of its activating cyclins had similar localization and could thus activate CDK1.

Two families of mitotic cyclins, A and B, associate with CDK1 to promote mitotic entry (reviewed in (54)). CDK1 tends to associate preferentially with CycA in early G2 and with CycB later. We first evaluated CycB. In uninfected cells, CycB was first detected as a dispersed cytoplasmic signal. At early prophase, it started to accumulate at the nuclear periphery and at the centrosomes. Later, CycB localized around condensed chromosomes and to the mitotic spindle. CycB signal was lost in anaphase, in agreement with its regulated degradation initiated by the activation of the anaphase promoting complex (APC) after the spindle assembly checkpoint is satisfied (**Suppl. Fig. 2A**). CycB localization in ZIKV-infected cells was unremarkable and indistinguishable from its localization in uninfected cells (**Suppl. Fig. 2B**). CycB accumulated and degraded through the cell cycle similarly in mock-or ZIKV-infected cells.

We next evaluated CycA. In mock infected hiNPC, CycA localized predominantly to the nucleus with dispersed cytoplasmic staining, as expected (96) (Fig. 6B). CycA signal was lost in late metaphase, in agreement with its known degradation initiated by APC. Starting 3h after ZIKV infection, however, infected cells started to display abnormally localized cytoplasmic CycA, often in the absence of any obvious nuclear signal (Fig. 6C, F). The number of cells showing this abnormal localization increased with time, and all ZIKV-infected cells had abnormal cytoplasmic CycA at 24 hpi, when CycA was particularly enriched around NS1. Similarly to CDK1, CycA and NS1 signals were intermixed (PCC, 0.58, Fig. 6C) rather than fully colocalized as those of PNKP and NS1 (Fig. 6D). All ZIKV-infected cells expressed detectable levels of CycA at 24 h post infection, as opposed to the expected degradation in a large subset of mock-infected cells (Fig. 6F). Our results thus indicate that ZIKV infection induces cytoplasmic accumulation of CycA and CDK1.

### ZIKV induces unscheduled activation of CycA/CDK1

ZIKV induced accumulation of cytoplasmic CycA and CDK1, presumably to trigger ER membrane rearrangements, correlated with gross mitotic abnormalities, and Rosco, which inhibits CDK1 inhibited the formation of RF. We thus asked whether the abnormal accumulation of CycA and CDK1 during ZIKV infection resulted in active CDK1 complexes at the sites of RF assembly.

We evaluated the kinase activity of CDK1 or CycA immunoprecipitates from 3 to 24 hpi. CDK1 activity increased in ZIKV infected hiNPC cells from the time of infection. Following an initial steady-state increase for up to 12 hpi, there was an exponential increase in CDK1 activity, reaching on average about 5-fold higher than in non-infected cells at 21 hpi (Fig. 7A). CDK1 kinase activity did not change much in asynchronous control cells in the 24 h, as expected (Fig. 7A, insert). Similarly, CycA antibody co-immunoprecipitated kinase activity was elevated in ZIKV-infected cells, to about 2-fold over control cells at 21 hpi, whereas there were no major changes in non-infected cells (Fig. 7A). Our results indicate that ZIKV induces unscheduled activation of cytoplasmic CycA/CDK1 activity, activation that continuously increases until 21-24 h post ZIKV infection.

**Figure 7.**
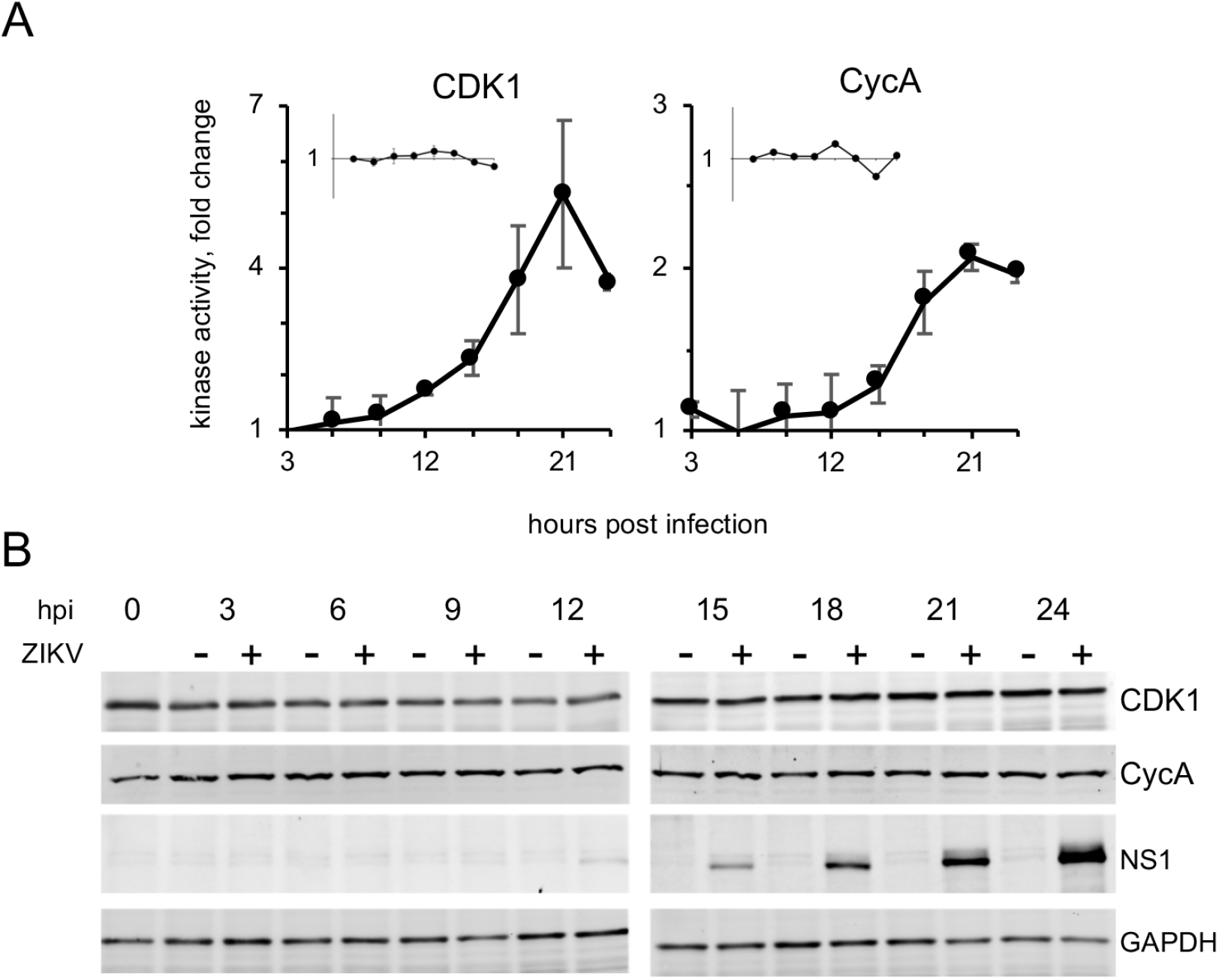
ZIKV infection induces unscheduled activation of cytoplasmic CycA/CDK1. (A) hiNPC were infected with ZIKV R1034, (MOI=2), or mock-infected and lysed at indicated times. CDK1 or CycA were immunoprecipitated and kinase activity was evaluated by immuno-kinase assays; fold change over mock-infected cells; inserts, mock-infected cells, fold change over T=0; n=2 (average +/- range). (B) hiNPC were infected with ZIKV R1034, (MOI=2), or mock infected, lysed at indicated times, and analyzed by Western blotting. Immuno-reactive bands were visualized by fluorescently labelled secondary antibodies. Images are representative of 3 independent experiments.

We next evaluated CycA and CDK1 levels through infection. CDK1 is constitutively expressed, and its level is constant during the cell cycle. CycA expression is tightly regulated and synchronized with cell cycle progression. Its accumulation begins in late G1, peaks in mid-S phase and declines in early mitosis (52, 53). In populations of unsynchronized uninfected cells, however, the average levels of CycA across the cell population are approximately constant, as different cells progress through the cell cycle at different times. No major differences in total CycA or CDK1 accumulation in the populations of ZIKV-infected cells were observed, and their levels were as constant as expected (Fig. 7B). Therefore, the elevated activity of CycA/CDK1 was not related to increased expression levels.

The elevated levels of CycA/CDK1 complexes in the cytoplasm thus result in increased CycA/CDK1 activity in infected cells. However, this elevated CycA/CDK1 activity is surprising in consideration of the abundant DNA damage (Fig. 3A, **Suppl. Fig. 1A**). We thus evaluated checkpoint activation.

### DNA damage-activated checkpoints are defective in ZIKV infection

As CDK1 was active in the presence of DNA damage, we tested checkpoint activation. The G2/M checkpoint inhibits CDK1 activity to prevent mitotic progression in the presence of damaged or incompletely replicated DNA. The unscheduled activation of CycA/CDK1 in the presence of DNA damage in ZIKV infection suggested dysfunctional checkpoint signaling. DNA damage activates the ATM and the ATR kinases, which in turn phosphorylate checkpoint kinases Chk1 and Chk2 to stop cell cycle progression (reviewed in (97)).

To test checkpoint signaling, we analyzed the activation of ATM, ATR, Chk1, and Chk2 with phospho-specific antibodies recognizing their activated phosphorylated forms (ATM, pS1981; ATR, pT1989; CHK1, pS345; CHK2, pT68). As expected, UV irradiation at 100 J/m^2^ resulted in the accumulation of the DNA damage marker γH2AX and activation of the damage sensing kinases ATM and ATR and checkpoint kinases Chk1 and Chk2, as detected by the presence of their specific phosphorylated forms at 1 h post irradiation (Fig. 8). There was also an accumulation of total γH2AX in ZIKV-infected cells, most obvious at 21 and 24 hpi. Unlike in UV-treated cells however, there was no obvious ATM activation in infected cells. The background levels of ATR activation in NPC were relatively high but did not differ between infected and mock-infected cells. This basal ATR activation in NPC likely results from their innate replication stress (98). As expected from the lack of activation of ATM or ATR, there was no downstream activation of Chk1 or Chk2 in ZIKV infected hiNPC either.

**Figure 8.**
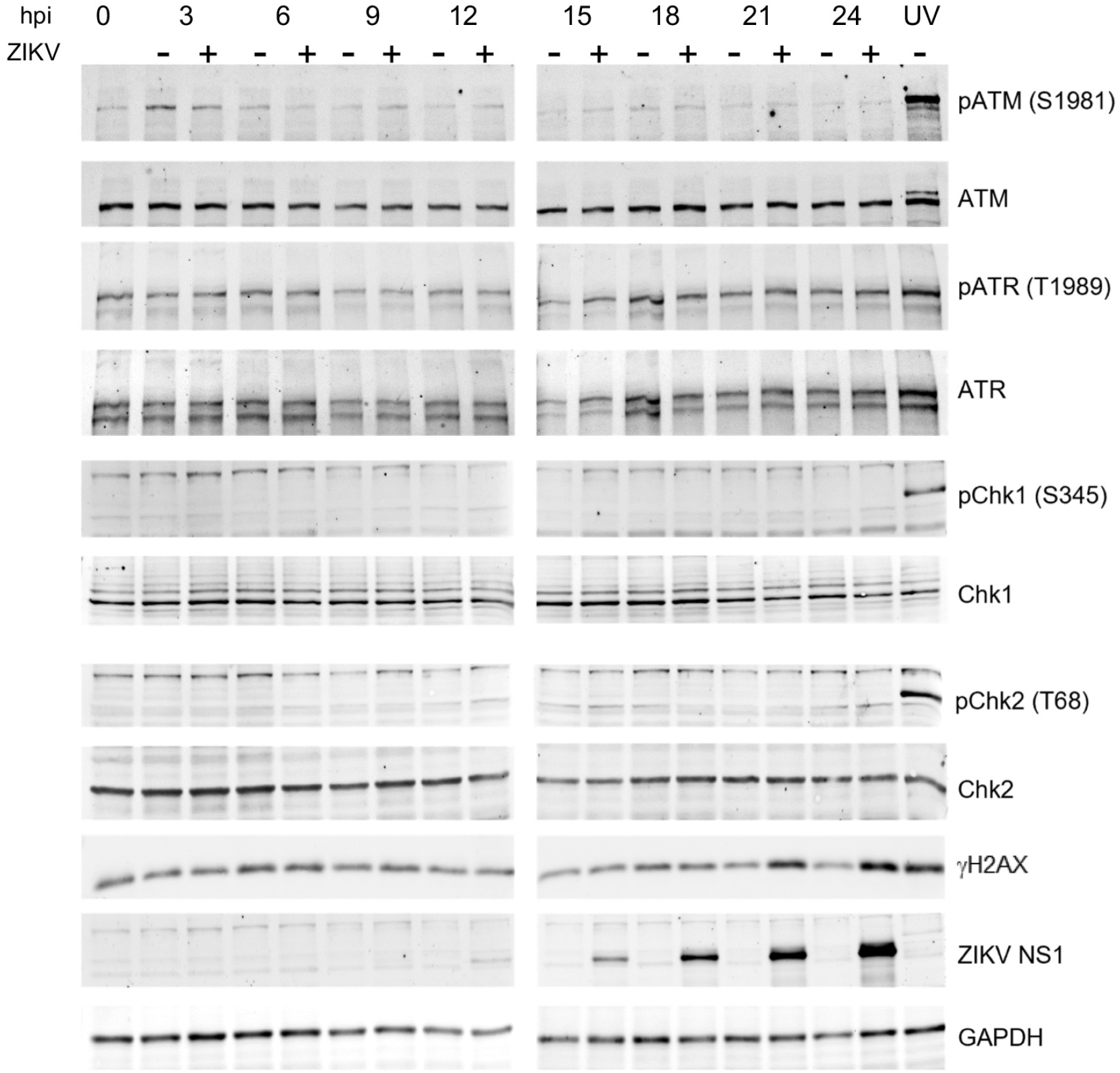
Neither G1/S nor G2/M DNA damage checkpoints are activated in ZIKV infected hiNPC. hiNPC were infected with ZIKV R1034 (MOI=2), or mock-infected, harvested at the indicated times, lysed, and analyzed for total and phosphorylated forms of checkpoint signaling (ATM, ATR) and effector (Chk1, Chk2) kinases by Western blot. DNA damage was induced in uninfected cells by UV irradiation (100 J/m^2^). Cells were allowed to recover for 1 h prior to harvesting. Immunoreactive bands were visualized by fluorescently labelled secondary antibodies. Images representative of 3 independent experiments.

In summary, ZIKV-infected cells failed to activate the cell cycle checkpoints despite the presence of DNA damage, CDK1 was thus activated and triggered mitosis in the presence of DNA damage, which remained unrepaired because of the functional PNKP depletion, thus resulting in MC.

## Discussion

Congenital ZIKV infections preferentially target fetal neural progenitor cells. Infection of these cells results in a series of neurodevelopmental abnormalities of which the most serious is microcephaly. In the present study, we show a link between ZIKV infection and the DNA damage repair enzyme PNKP, which is itself also linked to microcephaly. PNKP translocated to cytoplasmic clusters co-localizing with NS1, which is a marker of ZIKV RF. Two PNKP phosphatase inhibitors, A83B4C63 and A12B4C3, inhibited ZIKV replication in a dose-dependent manner and PNKP knockout HCT 116 cells were not permissive for ZIKV replication. ZIKV-infected human neural progenitors accumulated DNA damage but failed to activate the DNA damage checkpoint kinases Chk1 and Chk2 and CDK1 activity increased. Consequently, the cells underwent defective mitoses, producing grossly abnormal nuclear morphologies consistent with the phenotype of MC Fig. 9). ZIKV infection also induced cytoplasmic accumulation and activation of CycA/CDK1 complexes. Inhibition of cell division with Rosco, which inhibits CDK1 and CDK2, as well as other protein kinases not involved in mitosis like CDK5, inhibited the formation of ZIKV RFs and the production of ZIKV infectivity. Being an ATP competitor, the concentration of Rosco required to inhibit CDK is dependent on that of ATP. It has been shown that although 3 µM suffices to inhibit CDK at very low ATP concentrations, 100 µM is actually required to inhibit CDK at physiological ATP concentrations (about 0.5 µM or higher) (87). The well characterized specificity of Rosco against the human kinome has been described and discussed extensively many times before (83–87). Unscheduled activation of CycA/CDK1 and checkpoint failure thus lead to premature mitotic entry of infected cells in the presence of damaged or incompletely replicated DNA, which in turn lead to MC. It remains unclear at this time why nuclear depletion or cytoplasmic accumulation of PNKP would be required during the replication cycle of ZIKV, as are the mechanisms of PNKP recruitment to the RF, or of CycA/CDK1 to their proximity. It is also unclear why PNKP would participate in the replication of a cytoplasmic RNA virus, although it has a role in the replication of some nuclear DNA viruses (99, 100). We are in the process of addressing these yet outstanding issues.

**Figure 9.**
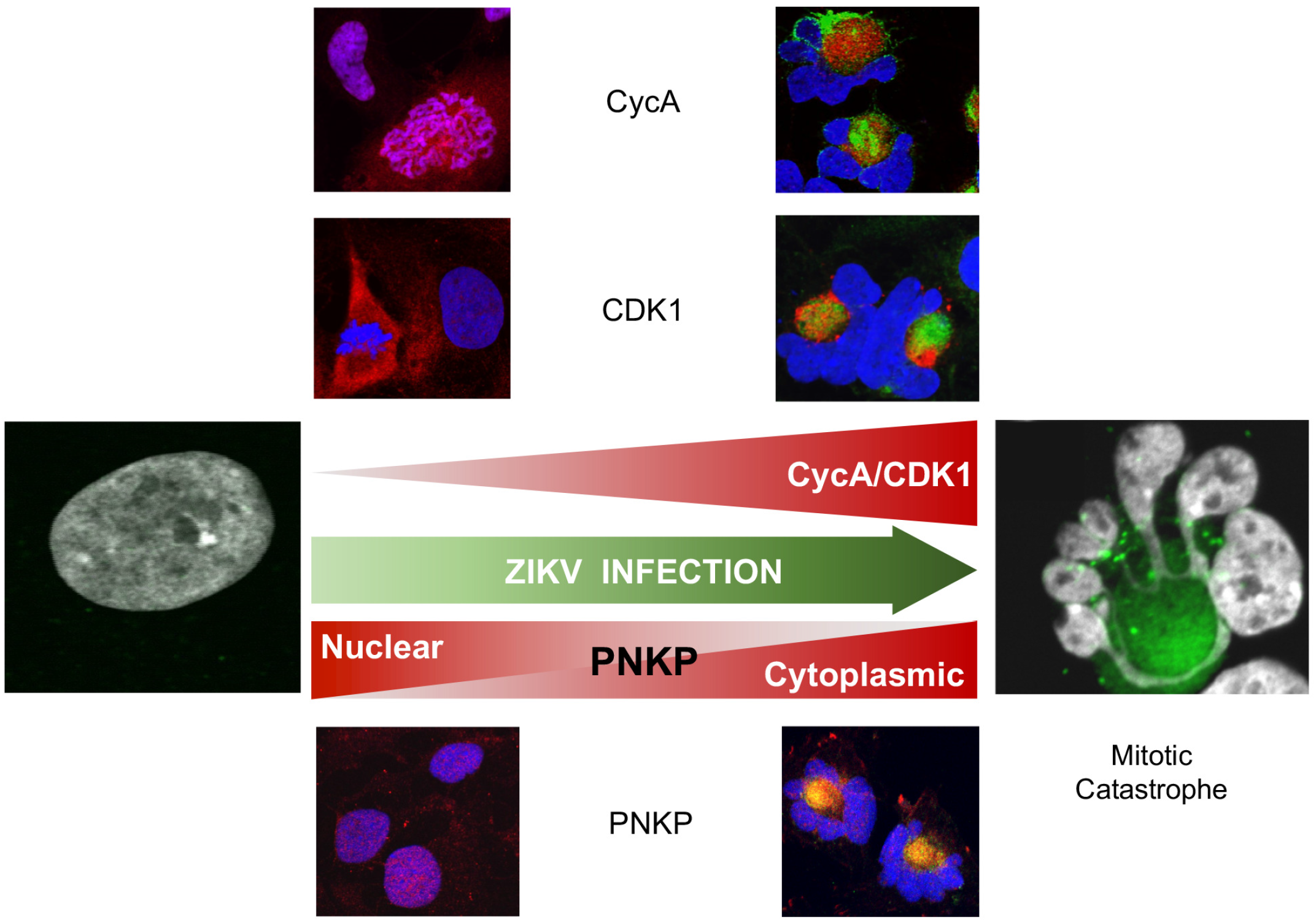
A model for the development of mitotic catastrophe in ZIKV-infected hiNPCs. ZIKV infection induces the activation of cytoplasmic CycA/CDK1 complexes, presumably to facilitate the formation of the viral replication factories, while sequestering PNKP away from the nucleus and into the replication factories. Unscheduled mitotic entry results in DNA damage, which cannot be repaired because of the functional depletion of nuclear PNKP activity; however, checkpoints are not activated and CDK1 activity continues to increase, resulting in mitotic catastrophe and subsequent neuronal progenitor cell death. Red, CycA, CDK1 or PNKP; green, NS1.

The PNKP gene (MIM605610, locus 19q13.33) was identified in 1999 (34, 35). It encodes a polynucleotide 5’-kinase 3′-phosphatase involved in DNA damage repair. PNKP is recruited to the sites of DNA damage to repair DNA breaks with so-called ‘dirty’ 3’-phosphate (3’-PO) and 5’-hydroxyl (5’-OH) termini. Restoring canonical DNA termini by the 5’-kinase and 3’-phosphatase enzymatic activities of PNKP is essential to repair DNA breaks bearing such ‘dirty’ termini. PNKP deficiencies hamper single and double strand break repair (44, 46, 47), increasing the frequency of spontaneous mutations and sensitizing cells to genotoxic stress (45). PNKP deficient cells accumulate DNA damage and chromosomal lesions, producing mitotic abnormalities including micronuclei and chromosomal bridges (44, 47).

Some PNKP genetic variants are associated with rare (less than 50 patients worldwide) autosomal recessive neurological disorders, presenting with neurodevelopmental or neurodegenerative symptoms. The most severe of these disorders produces early onset primary microcephaly with seizures and developmental delay (MCSZ, MIM 613402), also known as developmental and epileptic encephalopathy-10, DEE10) (31, 101). It is not entirely clear how different PNKP mutations contribute to either neurodevelopmental or neurodegenerative pathologies, but recent data indicate that residual levels of phosphatase or kinase activity may play a role. In patient-isolated fibroblasts, reduced phosphatase activity of PNKP mutants and inability to repair 3’-PO single stranded DNA breaks correlated with neurodevelopmental disorders (MCSZ), whereas reduced kinase activity in the context of functional phosphatase correlated with milder neurodegenerative pathologies (44).

Two PNKP phosphatase inhibitors, A83B4C63 and A12B4C3, inhibit ZIKV replication (Fig. 1A, B), at concentrations in the range of those required to inhibit other cellular outcomes of PNKP activity (1-50 μM -(66-68, 70-72) (Fig. 1C). Furthermore, PNKP knockout HCT 116 cells were not permissive for ZIKV replication, indicating that PNKP plays an important role in ZIKV replication. However, PNKP normally localizes to the nucleus and mitochondria (74, 102) and ZIKV replicates in cytoplasmic ER-derived RF (22, 24, 25). Surprisingly, PNKP clustered in the cytoplasm in ZIKV-infected cells co-localizing with NS1, a marker of RF (Fig. 2A, B, and C). As ZIKV infection did not affect total PNKP levels (Fig.1E), PNKP re-localizes to these cytoplasmic clusters from the nucleus, and possibly mitochondria. The resulting depletion of phosphorylated nuclear PNKP (Fig. 3B, **Suppl. Fig. 1C**) likely results in functional enzyme deficiency and the consequent accumulation of unrepaired DNA breaks (Fig. 3A, **Suppl. Fig. 1A**). A recently identified PNKP mutation in an individual with MCSZ (P101L) results in a novel nuclear export signal and consequently cytoplasmic PNKP localization (Jiang et al., unpublished data). Nuclear PNKP depletion, like the one induced by ZIKV, may therefore be sufficient to induce neurodevelopmental defects and microcephaly.

Following genotoxic stress, the DNA damage-activated ATM kinase phosphorylates PNKP at Ser114 to prevent PNKP proteasomal degradation, thus allowing efficient recruitment and accumulation of PNKP at the damage sites (73, 78, 79). Although ZIKV-infected cells accumulated DNA damage (Fig. 3A, Fig. 6, **Suppl. Fig. 1A**), there were no phosphorylated PNKP foci (Fig. 3B, **Suppl. Fig. 1C**). PNKP was thus not efficiently activated at DNA damage sites. PNKP is recruited to DNA breaks by direct interaction with the DDR scaffolding proteins, X-ray repair cross complementing protein 1 (XRCC1), or 4 (XRCC4) (76, 77, 103). PNKP could thus have been recruited to the viral RF by either. However, ZIKV-induced cytoplasmic PNKP localization was independent of XRCC1 or XRCC4, which both remained nuclear (Fig. 2E).

DNA damage, mitotic aberrations, accumulation of chromosomal abnormalities, and death of progeny cells after mitosis have all been reported in ZIKV-infected cells (104–110). Based on the analysis of the abundant and grossly abnormal nuclear morphologies in ZIKV-infected neural progenitors, seen by us (Fig. 2A, 3A, 4C, D) and others (104–106), we propose that infected NPC undergo MC. MC has been defined by “the nomenclature committee on cell death” as a “regulated oncosuppressive mechanism that impedes the proliferation or survival of cells unable to complete mitosis due to extensive DNA damage, problems with mitotic machinery, or failure of mitotic checkpoints” (111). Cells undergoing MC die at mitosis, slip out of mitosis and enter senescence, or proceed to interphase and die at subsequent cell cycles (82). MC is defined morphologically by distinct nuclear changes, including micronucleation (presence of micronuclei), multinucleation (presence of multilobular nuclei surrounded by nuclear envelope) or macronucleation (two or more nuclei collapsed into one and surrounded by nuclear envelope) (81). All the above MC phenotypes were present in ZIKV-infected NPC (Fig. 2A; 3A, B; 4C, D) and absent when mitosis was inhibited with Rosco (Fig. 5B, E).

Damaged DNA causes chromosome segregation failure during mitosis and subsequent defects in mitotic exit, due to abnormal, broken, linked, or entangled chromosomes. Experimental PNKP depletion produced mitotic abnormalities in cultured astrocytes (47). However, micronuclei and chromosomal bridges, indicative of chromosome segregation failure, were detected only in PNKP knockout and also irradiated cells, despite the presence of basal levels of DNA damage in the untreated cells. We and others have observed DNA damage, mitotic abnormalities (104–110) and MC during ZIKV infection in the absence of any exogenous genotoxic stress. ZIKV-induced MC would thus require abnormal mitotic entry in the presence of the damaged DNA. Here we show that ZIKV infection induced activation of cytoplasmic CycA/CDK1 (Fig. 7A), triggering unscheduled mitotic entry even in the presence of DNA damage, thus leading to MC. The activity of CDK1 was elevated throughout the infection, which prevents mitotic exit, resulting in MC.

Premature mitosis is deleterious and therefore entry into mitosis is highly regulated. DNA damage-regulated checkpoints respond to DDR signaling by arresting cells at the G2-M and G1-S transitions to allow DNA repair or induce regulated cell death or senescence if the damage is beyond repair (recently reviewed in (97)). The main signal transducers linking DNA damage responses to checkpoint effectors are the protein serine/threonine kinases ATM and ATR. Activated ATM and ATR initiate signaling to trigger the multiple pathways of DNA repair and activate the checkpoint kinases Chk1 and Chk2 to induce temporary cell cycle arrest at G1-S or G2-M. To this end, Chk1 and Chk2 inactivate the CDC25 family phosphatases, thus inhibiting mitotic cyclin/CDK1 and preventing mitotic entry (112).

Elevated CDK1 activity in the presence of DNA damage thus also indicates checkpoint failure. Indeed, ZIKV-infected cells failed to efficiently activate the checkpoint kinases Chk1 and Chk2, although they had about as much DNA damage (based on γH2AX levels) as uninfected cells treated with UV (Fig. 8). Defective ATR/Chk1 checkpoint has been reported previously in ZIKV infection in human NPC (110), while the ATM/Chk2 checkpoint was reported active, but only at 48 h post infection. In our studies, MC was induced between 12-15 h after infection and most of the infected cells presented a well-defined MC phenotype at 24 h, before the previously reported ATM and Chk2 activating phosphorylation. We analyzed checkpoint activation at early times, from 3 to 24 h post infection, which would be relevant for MC at 24 h. There was no activation of Chk1 or Chk2 in ZIKV infected hiNPC as compared to their levels of phosphorylation induced by UV irradiation (Fig. 8) despite the presence of abundant γH2AX staining in ZIKV infected hiNPC (Fig. 3A, **Suppl. Fig. 1A,** Fig. 8), and CDK1 was active (Fig. 7A), consistently with the lack of Chk1/2 activation during ongoing mitosis (113, 114).

The cytoplasmic CycA localization detected in ZIKV-infected hiNPC (Fig. 6C, F) is not entirely unusual. CycA localizes to the cytoplasm at the S/G2 transition where it initiates the activation of Plk1 through the phosphorylation of the cytoplasmic cofactor Bora (115). Plk1 activation in late G2 sets the commitment to mitosis (116). CDK1 and CycA accumulated in the cytoplasm of ZIKV-infected cells adjacent to ZIKV RF and all ZIKV-infected cells were positive for cytoplasmic CycA at 24 hours after infection (Fig. 6C, F). Thus, the unscheduled mitotic entry of ZIKV-infected hiNPC is likely triggered by this cytoplasmic accumulation of active CycA/CDK1 complexes.

Inhibition of cell duplication by inhibition of CDK1, CDK2 (and other kinases not involved in mitosis) with Rosco inhibited the formation of ZIKV RF (Fig. 5B), and the consequent production of cell-free infectious ZIKV virions (Fig. 5E). ZIKV RF are derived from modified rough ER (RER) membranes (22, 24–26). The morphology of ZIKV RF is predominantly tubular or cisternal, while the RER has predominantly sheet morphology, with the exception of the peripheral RER membranes that form the tubular networks. The ER undergoes sheet to tubule transformation during mitosis to enable the organelle partitioning to the daughter cells during cytokinesis (94, 95). CycA is involved in the reorganization of the ER network during mitosis in the *Drosophila* embryo (117) and in nuclear envelope breakdown, an event that initiates the ER rearrangement, in HeLa cells (118). We propose that ZIKV likely induces the activation and cytoplasmic accumulation of CycA/CDK1 to induce morphological changes to the ER membranes to facilitate the formation of the viral RFs. Curiously, CDKs continue to be shown to play important roles in the replication of RNA viruses whose replication requires no cell cycle progression (119).

Of the 25 genes linked to human autosomal recessive primary microcephaly (MCPH), 22 encode mitotic regulators (120). Mutations in the *MCPH2* gene, encoding the centrosomal protein Wdr62 linked to MCPH, delay mitotic progression due to spindle instability leading to spindle assembly checkpoint (SAC) activation, resulting in mitotic arrest and cell death (121). Mitotic delay in murine NPC, either by pharmacological intervention or in transgenic mouse models, results in premature differentiation or death of NPC and produces microcephaly in transgenic animals (122–124). Mitotic abnormalities in ZIKV-infected cells, including supernumerary centriole, multipolar spindle and chromosomal abnormalities, have been documented previously (104–106, 125). Here, we show that ZIKV infection caused the activation of cytoplasmic CycA/CDK1 and unscheduled mitotic entry in the presence of DNA breaks, which accumulated in infected cells due to the cytoplasmic localization of PNKP, leading to grossly abnormal mitoses culminating in MC.

Previous studies have proposed several alternative molecular mechanisms underlying ZIKV-induced microcephaly. The functions of several mitosis-associated proteins, CEP152 and pericentrin (centrosomal), CENPJ (centromere) and TBK1 kinase (centrosomal) are affected during ZIKV infection, leading to mitotic defects and premature differentiation or cell death (104, 125). Microcephaly-linked RNA binding protein musashi-1 (MSI1) has been proposed as a host factor promoting ZIKV replication (126). MSI1 binding to viral RNA limited binding to its endogenous targets, thus affecting the expression of proteins playing important roles in NPC function. ZIKV infection was also shown to inhibit the AKT-mTOR signaling pathway, which plays essential roles in brain development and autophagy, leading to defective neurogenesis (127). ZIKV NS4A also targets another microcephaly-related protein ANKLE-2 and thus disrupts neurogenic asymmetric cell division in neural stem cells leading to microcephaly in a drosophila model (128). Mounting evidence also supports a role for the immune responses in the fetal neuropathology of ZIKV infection (129–134). Congenital Zika virus syndrome is thus most likely a result of a combination of several mechanisms and systemic processes, which we are only beginning to understand.

Similar to defects in mitotic genes, genetic defects affecting DNA damage responses often associate with neurodevelopmental defects presenting with microcephaly (135). We propose that the functional PNKP deficiency and dysregulation of CycA/CDK1 mitotic activity during ZIKV infection of neural progenitors result in MC, contributing to the molecular mechanism underlying neurodevelopmental pathologies of congenital ZIKV infections.

## Acknowledgments

These studies were funded by the NIH National Institute of Neurological Disorders and Stroke (5R21NS111416) and Canadian Institutes of Health Research (PJT168869). LMS is a BWF investigator in the pathogenesis of infectious disease. The authors acknowledge the support from the Baker Institute and the College of Veterinary Medicine, Cornell University. The authors thank Dr. Roy Golsteyn (University of Lethbridge, Canada) for his advice and insightful comments regarding mitotic catastrophe and Dr. Dennis Hall (University of Alberta, Canada) for providing the PNKP inhibitors A12B4C3 and A83B4C63.

